# Cellular and widefield imaging of sound frequency organization in primary and higher-order fields of the mouse auditory cortex

**DOI:** 10.1101/663021

**Authors:** Sandra Romero, Ariel E. Hight, Kameron K. Clayton, Jennifer Resnik, Ross S. Williamson, Kenneth E Hancock, Daniel B Polley

## Abstract

The mouse auditory cortex (ACtx) contains two core fields – A1 and AAF – arranged in a mirror reversal tonotopic gradient. The best frequency (BF) organization and naming scheme for additional higher-order fields remain a matter of debate, as does the correspondence between smoothly varying global tonotopy and heterogeneity in local cellular tuning. Here, we performed chronic widefield and 2-photon calcium imaging from the ACtx of awake *Thy1*-GCaMP6s reporter mice. Data-driven parcellation of widefield maps identified five fields, including a previously unidentified area at the ventral posterior extreme of the ACtx (VPAF) and a tonotopically organized suprarhinal auditory field (SRAF) that extended laterally as far as ectorhinal cortex. Widefield maps were stable over time, where single pixel BFs fluctuated by less than 0.5 octaves during a one-month imaging period. After accounting for neuropil signal and frequency tuning strength, BF organization in neighboring layer 2/3 neurons was intermediate to the heterogeneous salt and pepper organization and the highly precise local organization that have each been described in prior studies. Multiscale imaging data suggest there is no ultrasonic field or secondary auditory cortex in the mouse. Instead, VPAF and a dorsoposterior field (DP) emerged as the strongest candidates for higher-order auditory areas.

## Introduction

Topographic connections between the sensory receptor epithelia and downstream brain nuclei form well in advance of sensory experience and reflect the patterning of molecular guidance cues (Rakic et al. 2009; Cramer and Gabriele 2014). In the rodent auditory system, topographic connections linking the medial geniculate body (MGB) of the thalamus and the auditory cortex (ACtx) form in the last week of embryonic development, approximately one week prior to the onset of spontaneously generated intrinsic signaling between the ear and the brain, and 2 weeks prior to ear canal opening and the onset of hearing (Gurung and Fritzsch 2004; Tritsch et al. 2007; Polley et al. 2013). Cortical maps are not organized into a single, topographic gradient, but rather as a mosaic of repeating gradients, separated from one another by “mirror reversals” in receptor epithelium mapping. These individual fields of the ACtx can exhibit specializations in functional processing that reflect different origins of thalamic inputs as well as regional variations in the source, though not precision, of local and long-range intracortical connections (Rose and Woolsey 1949a; Lee and Winer 2005; Winer et al. 2005). Core fields of the ACtx receive their predominant thalamic input from ventral division of the MGB (MGBv), which confers well-defined frequency tuning arranged into smoothly varying tonotopic gradients (Merzenich and Brugge 1973; Andersen et al. 1980; Winer et al. 2005; Hackett 2011). Higher-order ACtx fields are innervated by non-primary divisions of the MGB and from intracortical inputs originating outside of the auditory cortex (Andersen et al. 1980; Reale and Imig 1980; Schreiner and Cynader 1984; Lee and Winer 2005; Winer et al. 2005; Hackett 2011). Higher-order cortical areas show stronger selectivity for processing conspecific communication sounds (Schneider and Woolley 2013; Norman-Haignere et al. 2015), greater involvement in cross-modal plasticity (Lomber et al. 2010) and stronger state-dependent modulation by cognitive influences such as task demands and learning (Dong et al. 2013; Atiani et al. 2014; Elgueda et al. 2019).

The mouse is among the most popular model systems for studies of cortical sound processing and plasticity, but fundamental aspects of core and higher-order cortical field organization remain unclear. The mirror reversal in tonotopy between the primary auditory cortex (A1) and an anterior auditory field (AAF), which has been observed in dozens of species (Kaas 2011), has been questioned in the mouse where instead several groups have described a region at the border of A1 and AAF without well-defined selectivity for pure tones (Issa et al. 2014, 2016; Liu et al. 2019). The systematic mapping of preferred frequency in core fields is well-accepted at mesoscale resolution (Stiebler et al. 1997; Hackett et al. 2011; Guo et al. 2012), but remains a point of contention at the cellular scale, with some reports describing a heterogeneous salt and pepper organization, while others describe a precise relationship at all spatial scales (Bandyopadhyay et al. 2010; Rothschild et al. 2010; Winkowski and Kanold 2013; Issa et al. 2014; Panniello et al. 2018; Liu et al. 2019; Tischbirek et al. 2019). The secondary field, A2, was named without having established that its primary thalamic input arose from the higher-order subdivisions of the MGB (Stiebler et al. 1997). Instead, recent evidence reports that A2 receives its primary input from the MGBv, with minimal input from the higher-order dorsal subdivision, the MGBd, raising questions about whether there are any fields in the mouse ACtx that are appropriately described as higher-order and, if so, where they are located (Ohga et al. 2018). Even the name and location of A1 is not beyond dispute, where some groups refer to it instead as a “dorsomedial field” (Tsukano, Horie, Ohga, et al. 2017).

Confusion about the basic features of mouse ACtx organization stems to no small extent from differences in methodology. Traditional microelectrode mapping cannot reveal local organization at a cellular scale and cannot easily be performed in awake animals. Imaging of intrinsic hemoglobin or flavoprotein fluorescence signals can be used to visualize the entire ACtx at once, but provide a less-direct, even lower resolution map than multiunit microelectrode recordings (Kalatsky et al. 2005; Tsukano, Horie, Ohga, et al. 2017; Tischbirek et al. 2019). 2-photon imaging of calcium signals through bulk-loaded dyes or virus mediated gene transfer can be performed in awake mice and provides direct access to a cellular signal closely linked to spiking, but does not easily provide the even, stable expression over large areas needed to image multiple fields of the ACtx (Bandyopadhyay et al. 2010; Rothschild et al. 2010; Kato et al. 2016; Kuchibhotla et al. 2017; Francis et al. 2018; Tischbirek et al. 2019). A promising approach to resolve the disputed organization of the mouse ACtx at multiple scales comes from imaging of awake transgenic mice that express the genetically encoded calcium indicator GCaMP6 in select cell types (Issa et al. 2014, 2016; Babola et al. 2018; Panniello et al. 2018; Liu et al. 2019). Here, we performed multiscale imaging from excitatory neurons in the ACtx of *Thy1*-GCaMP6s reporter mice to delineate mesoscale map organization through widefield, epifluorescence imaging and cellular organization of frequency selectivity through 2-photon imaging.

## Methods

### Experimental model and subject details

Adult mice of either sex were used for all experiments in the study. All procedures were approved by the Massachusetts Eye and Ear Infirmary Animal Care and Use Committee and followed the guidelines established by the National Institute of Health for the care and use of laboratory animals. *Thy1*-GCaMP6s mice (Jackson labs stock number 025776) were used in a subset of auditory brainstem response (ABR) measurements. Imaging experiments were performed on male and female offspring of *Thy1*-GCaMP6 crossed with CBA/CaJ, a strain that retains good ABR thresholds through adulthood (Zheng et al. 1999). Mice were housed in group cages until cranial window implantation, at which point they were housed individually. Mice were maintained in a 12h light/dark cycle, with food and water available *ad libitum*. Mice were between 5-7 weeks at the time of cranial window surgery and were no older than 20 weeks by the time the last imaging session was completed.

### Auditory brainstem response measurements

Mice were anesthetized with ketamine and xylazine (100/10 mg/kg for ketamine/xylazine, respectively, with boosters of 50 mg/kg ketamine given as needed). Core body temperature was maintained at 36.5° with a homeothermic blanket system. Acoustic stimuli were presented via in-ear acoustic assemblies consisting of two miniature dynamic earphones (CUI CDMG15008–03A) and an electret condenser microphone (Knowles FG-23339-PO7) coupled to a probe tube. Stimuli were calibrated in the ear canal in each mouse before recording.

ABR stimuli were tone bursts (4-64 kHz in 0.5 octave increments), 5 ms duration with a 0.5 ms rise-fall time delivered at 27 Hz, and alternated in polarity. Intensity was incremented in 5 dB steps, from 20-80 dB SPL, or as high as 100 dB SPL in cases with elevated thresholds. ABRs were measured with subdermal needle electrodes positioned beneath both pinna (+ and −) and the base of the tail (ground). Responses were amplified (gain = 10,000), filtered (0.3–3 kHz), and averaged (1024 repeats per level). ABR threshold was defined as the lowest stimulus level at which a repeatable wave 1 could be identified. ABR Recordings were made from *Thy1*-GCaMP6s mice at three ages: 7-8 weeks (9 ears, 5 mice), 14 weeks (10 ears, 5 mice) and 20 weeks (8 ears, 4 mice). ABR recordings were made from *Thy1*-GCaMP6s × CBA mice at the same ages: 7-8 weeks (11 ears, 6 mice), 14 weeks (6 ears, 3 mice) and 20 weeks (6 ears, 3 mice).

### Preparation for chronic imaging

Glass cover slips were first etched in piranha solution (H2O2 mixed with H2SO4 in a 3:1 ratio) and stored in 70% ethanol. A 4mm diameter cover slip was centered and affixed to a pair of 3mm cover slips (#1 thickness, Warner Instruments) using a transparent, UV-cured adhesive (Norland Products). Windows were stored in double deionized water and rinsed with sterile saline before use. On the day of surgery, animals were anesthetized with isoflurane in oxygen (5% induction; 1.5-2% maintenance). After removing the periosteum from the dorsal surface of the skull, an etchant (C&B Metabond) was applied for 30 sec to create a better adhesive surface. Custom stainless-steel head fixation hardware (iMaterialise) was bonded to the dorsal surface of the skull with dental cement (C&B Metabond) mixed with India ink. A 3mm circular craniotomy was made atop the ACtx with the combination of a scalpel and a pneumatic dental drill with diamond burr (head diameter 1/10 mm, NeoDiamond – Microcopy Dental). The coverslip was then lowered into place using a 3-D manipulator and bonded to the surrounding regions of the skull to create a hermetic seal. Post-operative injections of Buprenex (0.05 mg/kg) and Meloxicam (0.1 mg/kg) were administered and the mice were allowed to recover in a warmed chamber. Imaging began 5-7 days later.

### In vivo calcium imaging

Widefield epifluorescence images were acquired with a tandem-lens microscope (THT-microscope, SciMedia) configured with low-magnification, high-numerical aperture lenses (PLAN APO, Leica, 2× and 1× for the objective and condensing lenses, respectively). Blue illumination was provided by a light-emitting diode (465 nm, LEX2-LZ4, SciMedia). Green fluorescence passed through a filter cube and was captured at 20Hz with a sCMOS camera (Zyla 4.2, Andor Technology). Two-photon excitation was provided by a Mai-Tai eHP DS Ti:Sapphire-pulsed laser tuned to 940 nm (Spectra-Physics). The beam spot size was controlled with variable expander optics (Thorlabs) and the intensity was adjusted with a variable attenuator (Thorlabs) and Pockels cells (Conoptics).

Imaging was performed with a 16×/0.8NA water-immersion objective (Nikon) from a 512 × 512 pixel field of view at 30Hz with a Bergamo II Galvo-Resonant 8 kHz scanning microscope (Thorlabs). Scanning software (Thorlabs) was synchronized to the stimulus generation hardware (National Instruments) with digital pulse trains. Widefield and 2-photon microscopes were rotated by 50-60 degrees off the vertical axis to obtain images from the lateral aspect of the mouse cortex while the animal was maintained in an upright head position. Imaging was performed in light-tight, sound attenuating chambers (N=12 mice for widefield imaging and N=4 mice for 2-photon imaging). Animals were monitored during the experiment with modified cameras (PlayStation Eye, Sony) coupled to infrared light sources. For widefield imaging, the focal plane was set to be approximately 200 μm below the pial surface. For 2-photon imaging, the imaging depth ranged from 175-225 μm below the pial surface, in layer 2/3.

### Auditory stimulation for imaging experiments

Stimuli were generated with a 24-bit digital-to-analog converter (National Instruments model PXI-4461) using scripts programmed in MATLAB (MathWorks) and LabVIEW (National Instruments). Stimuli were presented via free-field tweeters positioned 10 cm (2-photon system) or 25 cm (widefield system) from the left (contralateral) ear. Free-field stimuli were calibrated before recording using a wide-band ultrasonic acoustic sensor (Knowles Acoustics, model SPM0204UD5). Pure tones were pseudorandomly presented at variable frequencies (4-64 kHz in 0.5 octave steps) and intensities (0-70 dB SPL in 10 dB steps) such that each of the 72 unique frequency-intensity combinations were presented 20 times each. Tone duration was 50 ms. Trial length was either 3s (2-photon imaging) or 3.5s (widefield imaging).

### Image processing – Widefield

Raw data was first downsampled from the native 1200 × 1200 pixel resolution to 256 × 256 pixels. Slow drifts in the fluorescence signal were removed from the measurement by concatenating all frames for an individual imaging session and computing a temporal baseline (F0) for each pixel from the linear fit of a 10s sliding window incremented in 5s steps (Chronux toolbox, Matlab). The fractional change in fluorescence was defined for each frame as a percent change in signal from the temporally detrended average signals as (ΔF/F0) × 100. Individual trials were then averaged across the 20 repetitions and temporally filtered with a GCaMP6s impulse response deconvolution kernel (Chen et al. 2013). Fluorescence data were spatially denoised with Lucy-Richardson deconvolution using a Gaussian filter (470 μm width). For each stimulus, the temporal peak in the sound-evoked response period was defined independently for each pixel as the frame with the maximum percent change within a 0.75s period following stimulus onset, which was then averaged with the immediately preceding and following frame. For each pixel, baseline activity levels were defined by creating a histogram of percent change amplitudes during the 0.5s pre-stimulus period (25 frames × 72 stimuli). The response amplitude for each tone/level combination was then expressed in units of standard deviations (z-score) relative to the distribution of baseline activity levels.

### Image processing – two-photon

Imaging data were processed with Suite2p, a publicly available software package that provides a complete pipeline for processing calcium-dependent fluorescence signals collected with 2-photon microscopes (Pachitariu et al. 2016; Stringer and Pachitariu 2019). Briefly, fluorescence data were collected at 2x digital magnification and processed in four stages:

#### Frame registration

Brain movement artifacts are removed through a phase correlation process that estimates the XY offset values that bring all frames of the calcium movie into register. Suite2p emphasizes correcting for movement artifacts at high spatial frequencies by first applying spatial whitening before computing a cross-correlation map. A non-rigid method is then used for phase correction that divides the movie into independent blocks and computes the optimal XY offset for each discrete segment before applying the interpolated pixel shift function to the original image.

#### Detecting regions of interest (ROI)

Suite2p then identifies candidate cellular ROIs using a generative model with three key terms: *i)* a model of ROI activity, *ii)* a set of spatially-localized basis functions to model a neuropil signal that varies more gradually across space, and *iii)* Gaussian measurement noise. Fitting of this model to data involves repeatedly iterating stages of ROI detection, activity extraction, and subsequent pixel re-assignment.

#### Signal extraction and spike deconvolution

Suite2p then extracts a single fluorescence signal for each ROI by modelling the uncorrected fluorescence as the sum of three terms: *i)* a somatic signal due to an underlying spike train, *ii)* a neuropil trace scaled by an ROI-specific coefficient, and *iii)* Gaussian noise (Stringer and Pachitariu 2019). The uncorrected fluorescence is first extracted by averaging all signals within each ROI. The neuropil trace is then computed as the average signal within an annular ring surrounding each ROI. The neuropil component is different from those identified during ROI detection, which implicitly uses pixels inside ROIs, and are not scaled by a contamination factor. Neuropil scaling coefficients and somatic fluorescence are then simultaneously estimated using an unconstrained nonnegative deconvolution, using exponential kernels.

#### Cellular identification

With a fluorescence trace assigned to each identified ROI, the final stage in the Suite2p pipeline involves identifying the subset of ROIs that correspond to neural somata. Suite2p utilizes a semi-automated approach by first labelling ROIs as cells or non-cells based on various activity-dependent statistics, before a final manual curation step.

#### Response amplitude calculation

For the sake of direct comparison to widefield imaging, we computed the response amplitude of the 2-photon signal before and after neuropil correction using the same approach of identifying the peak response period for each stimulus and expressing this value as a z-score relative to the distribution of pre-stimulus baseline values.

### Registering images across sessions

To compare separate widefield imaging sessions from the same mouse, we first obtained images of the vasculature from the mean of the raw image stack. An Affine transformation matrix (‘*imregtform*’ Matlab function) was then computed between any pair of imaging sessions. The optimal Affine transformation matrix to align two images was identified using gradient descent to minimize the mean squares difference between the two images, within a maximum of 10,000 iterations.

To register the 2-photon and widefield images, an image of the surface vasculature was first obtained through the 16x objective using a CCD camera under epifluorescent illumination. We identified a set of five vascular landmarks contained in both the reference image collected on the tandem-lens widefield microscope and the target image collected through the 16x objective and the CCD camera on the 2-photon microscope. Pairs of points from the images were used to compute an affine transformation matrix and optimally align surface vasculature landmarks collected with the two imaging systems. The transformation matrix was then applied to the image acquired from the galvo-resonant scanner under 2-photon excitation. All 2-photon imaging sessions for a given mouse were registered to a single reference widefield imaging session.

### Data analysis

Except for the specific analysis of map changes over time, all analyses were performed only on the first widefield imaging session from each mouse. This way, no single mouse contributed more data than any other and there was no bias in selecting any particular type of imaging data.

#### Response threshold estimation

The minimum response threshold was estimated independently for each individual pixel in the widefield image or cell in the 2-photon image. Threshold was operationally defined as the lowest sound intensity for which the response to two adjacent tone frequencies were at least two standard deviations above the distribution of pre-stimulus baseline values.

#### Best frequency (BF) estimation

Frequency response functions were obtained by averaging the response at threshold, 10 dB above threshold and 20 dB above threshold. The BF was defined from the weighted sum of the responses for each of the test frequencies on an octave-based scale. Only pixels with BF response amplitudes with z-score values ≥ 2 were used for subsequent analyses.

#### Frequency tuning bandwidth

Tuning widths for each pixel was determined from the range of frequencies with response amplitudes z-scores ≥ 2 at 10dB above threshold.

#### Strength of tonotopy

For each pair of pixels, i and j, located at cortical positions pi and pj, respectively, a BF gradient vector was defined as the BF at site i minus the BF at site j, normalized by the Euclidean distance between pi and pj, all multiplied by a unit vector in the direction from pi to pj as:

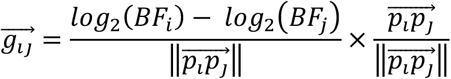

The resulting gradient, 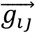, points from the pixel with the lower BF to the site with the higher BF and has a length proportional to the size of the change in BF normalized by the physical separation of the pixels. For each pixel *i*, a “tonotopic vector” was defined as the vector average of all the gradients between it and all the other pixels in the same field as:

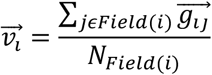

where *Field(i)* is the collection of pixels that belong to the same cortical field as pixel *i*, and *NField(i)* is the number of pixels in this field. The vector strength was calculated for each auditory field and defined as the magnitude of the vector average of all the tonotopic vectors that belong to a given cortical field:

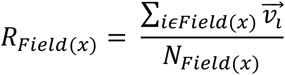

where Field(x) corresponds to all the pixels that belong to the auditory field x, NField(x) is the number of pixels in this field, and *R_Field(x)_* is the vector strength of the given field.

#### Similarity index

Modules with similar frequency tuning bandwidth or response threshold were identified by thresholding the BW10 and threshold maps at the highest and lowest quartile values and identifying the regional maxima or minima of 4×4 connected neighborhood of pixels with a minimum distance to another peak of 0.25 mm. Radial vectors were drawn from each and a Similarity Index (SI) between the center and the pixels falling along each vector was computed as:

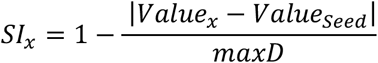

where *valuePointx* is the value of the response property (BW10 or ThrBF) at a pixel falling on the radial vector, valueSeed is the value of the response property (BW10 or ThrBF) at the seed pixel; and *maxD* is the maximal possible difference in the response property across the map. To compute the similarity that would occur by chance, the same procedure was repeated after shuffling the positions of the pixels 10,0 times. Module size was determined as the radial distance at which the actual SI values intersected with the mean of the shuffled SI values.

#### Strength of frequency tuning

To determine the strength of frequency tuning for each pixel (widefield) or cell (2-photon), we first identified the frequency/level combination from the entire frequency response area (FRA) with the highest response amplitude. We then determined the response amplitude for the adjacent frequencies and levels and calculated the average response amplitude from these five points. Of the remaining 67 frequency-level combinations, we selected five points at random and calculated the sensitivity index, d-prime (d’), to reflect the difference between the response amplitudes near the preferred stimulus versus stimuli selected at random. The process of selecting five random points was repeated 1000 times and the average d’ was operationally defined as the tuning quality.

#### Parcellation of auditory fields

We adapted the standard approach of defining the boundary between two adjacent fields according to reversals or abrupt shifts in the mapping of the receptor epithelia onto the cortical surface. We first identified the center of the four low-frequency points in A1, AAF, SRAF and VPAF (BF < 16 kHz). From each of these four low-frequency hubs, a set of 1440 radial vectors were drawn, at angles ranging from 0 to 360 degrees (step size = 0.25°). The mean BFs were calculated along each radial vector (± 1°). BFs were projected along each radial axis and fit to a smoothing spline (‘*fit*’ Matlab function). The reversal was defined as the point at which the first maxima was detected along the smoothed profile. If a reversal was not detected (for example at the map edges), the field boundary was drawn at the point where pixels were no longer sound-responsive according to the criteria above. To identify the boundary of the dorsal posterior field (DP), we computed the local BF gradients within A1 and created an XY map of the vector angle. The boundary between A1 and DP was aligned with the spatial shift in the BF vector phase map.

#### Widefield versus 2-photon frequency tuning

To relate the frequency tuning preference for individual neurons measured during 2-photon imaging to the underlying frequency selectivity in the widefield map, we first re-scaled the downsampled 256×256 widefield pixel map back to the native 1200×1200 pixel map. We then identified the individual pixels that correspond to the area of the neural ROI identified in Suite2p. The difference in the BF between the somatic ROI and the matching pixels of the widefield map was calculated as:

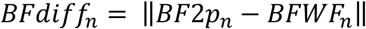

Where *BF2p_n_* is the BF from the 2-photon session and the *BFWF_n_* is the mean BF from the corresponding widefield ROI. The BF from the somatic ROI in the 2-photon session was calculated before and after the neuropil correction was applied.

#### Local BF heterogeneity

Variation in local BF tuning was measured from neuropil-corrected 2-photon imaging data. Within a given field of view, all somatic ROIs were identified within a 50 μm radius of the reference cell. Provided that a minimum of 5 cells were identified within this area, the median BF was computed across all cells within this local neighborhood. The absolute value of the BF difference for each cell versus the neighborhood median was calculated before repeating the process with a different reference neuron. The interquartile range of this BF distribution was operationally defined as the local BF heterogeneity.

#### Global tonotopic organization

To quantify the precision of global tonotopic organization from heterogeneous local cellular frequency tuning, we projected a radial vector from the center of the low-frequency hub to the high-frequency end of the tonotopic gradient. Position along the vector was normalized from 0-1 (corresponding to the low- and high-frequency extremes of the BF vector respectively). We then identified the global tonotopic vector that passed most directly through the center of each individual field of view from a single 2-photon imaging session. Each cell was then projected onto the nearest point of this global tonotopic vector. The Pearson correlation was defined by the BF of each neuropil-corrected cell and its position on the global tonotopic vector. Confidence intervals for the local BF heterogeneity and global tonotopic correlation coefficient were calculated by bootstrapping (10,000 iterations).

### Histology

A subset of mice (N=3) were anesthetized with isoflurane (5% in oxygen induction, 1.5% maintenance) after the final imaging session and the cranial window was removed. Points along the medial, lateral, rostral and caudal edges of the tonotopically organized areas were identified relative to surface vascular landmarks. A silicon probe (NeuroNexus) was mounted on a 3-D positioner and was dipped in (for 10s) and out (for 10s) ten times into red fluorescent dye (3 mg Di-I [1,1’-dioctadecyl-3,3,3’,3’-tetramethylindocarbocyanine perchlorate] per 100 μL acetone; Sigma-Aldrich, St. Louis, MO). The probe was inserted several hundred microns into the cortex at each of the four designated points and left in place for 30 minutes, re-applying the Di-I in between each placement. Following the last insertion, animals were prepared for transcardial perfusion with 4% paraformaldehyde in 0.01M phosphate buffered saline. Brains were extracted and stored in 4% paraformaldehyde for 12 hours before transferring to cryoprotectant (30% sucrose) for 48 hours. Sections (40 μm) were cut using a cryostat (Leica CM3050S), mounted on glass slides, coverslipped (Vectashield with DAPI), and imaged with an epifluorescence microscope (Leica).

### Statistical analysis

All statistical analysis was performed with Matlab (Mathworks). Descriptive statistics are reported as mean ± SEM unless otherwise indicated. In cases where the same data sample was used for multiple comparisons, we used the Holm-Bonferroni correction to adjust for the increased probability of Type-I error. Non-parametric statistical tests were used in cases where data samples did not meet the assumptions of parametric statistical tests. The relationship between FRA d’ and tuning heterogeneity was quantified with a permutation test iterated 500,000 times per auditory field. For any given iteration, each cell was re-assigned a d’ value at random, ensuring that the sample of d’ values was equivalent to the true distribution of d’ values. The outcome measures of interest, either the BF heterogeneity interquartile range or the global tonotopic correlation were then recomputed using the permuted data set and the linear relationship between the outcome measures of interest and the permuted d’ values were quantified for each iteration of the permuted and actual datasets with the Pearson R correlation coefficient. For plotting purposes, confidence intervals were computed as the standard error of the actual and permuted distributions. Statistical significance was established by determining the proportion of sampled permutations that exceeded the Pearson R of the true dataset. Statistical significance was defined as P < 0.05 for all tests.

## Results

### A transgenic mouse model for widefield calcium imaging that retains good hearing

We began our imaging studies in *Thy1-GCaMP6s* mice, which were developed for 2-photon imaging of the ultrasensitive genetically encoded calcium indicator GCaMP6s in cortical pyramidal neurons (Chen, Cichon, et al. 2012; Dana et al. 2014). We observed that these mice develop a substantial high-frequency hearing loss as early as 7 weeks postnatal, as indexed by wave 1 of the auditory brainstem response (ABR) (**Fig. 1A-B**). To circumvent this problem, we crossed the *Thy1*-GCaMP6s line to the CBA/CaJ line, which retains excellent hearing into adulthood, and confirmed that high-frequency ABR thresholds in *Thy1*-GCaMP6s × CBA mice were lower than the *Thy1*-GCaMP6s at all ages (Wilcoxon Rank-Sum, p < 0.0005 for all ages) and did not change through 20 weeks postnatal in the crossed strain (Kruskal-Wallis, *Thy1*-GCaMP6s × *CBA*, p = 0.38; *Thy1*-GCaMP6s, p < 0.005). GCaMP expression in the *Thy1*-GCaMP6s × *CBA* mouse was observed in the cytoplasm and neuropil of neurons throughout the cortical column (**Fig. 1C**). Once we were satisfied with the peripheral hearing status and cortical expression levels in *Thy1*-GCaMP6s × CBA mice, we manufactured custom head restraint hardware that left the ears unobstructed and adapted a surgical approach to implant a glass coverslip over the lateral areas of the skull so that the full extent of the auditory fields could be visualized (**Fig. 1D**) (Goldey et al. 2014).

**Figure 1.**
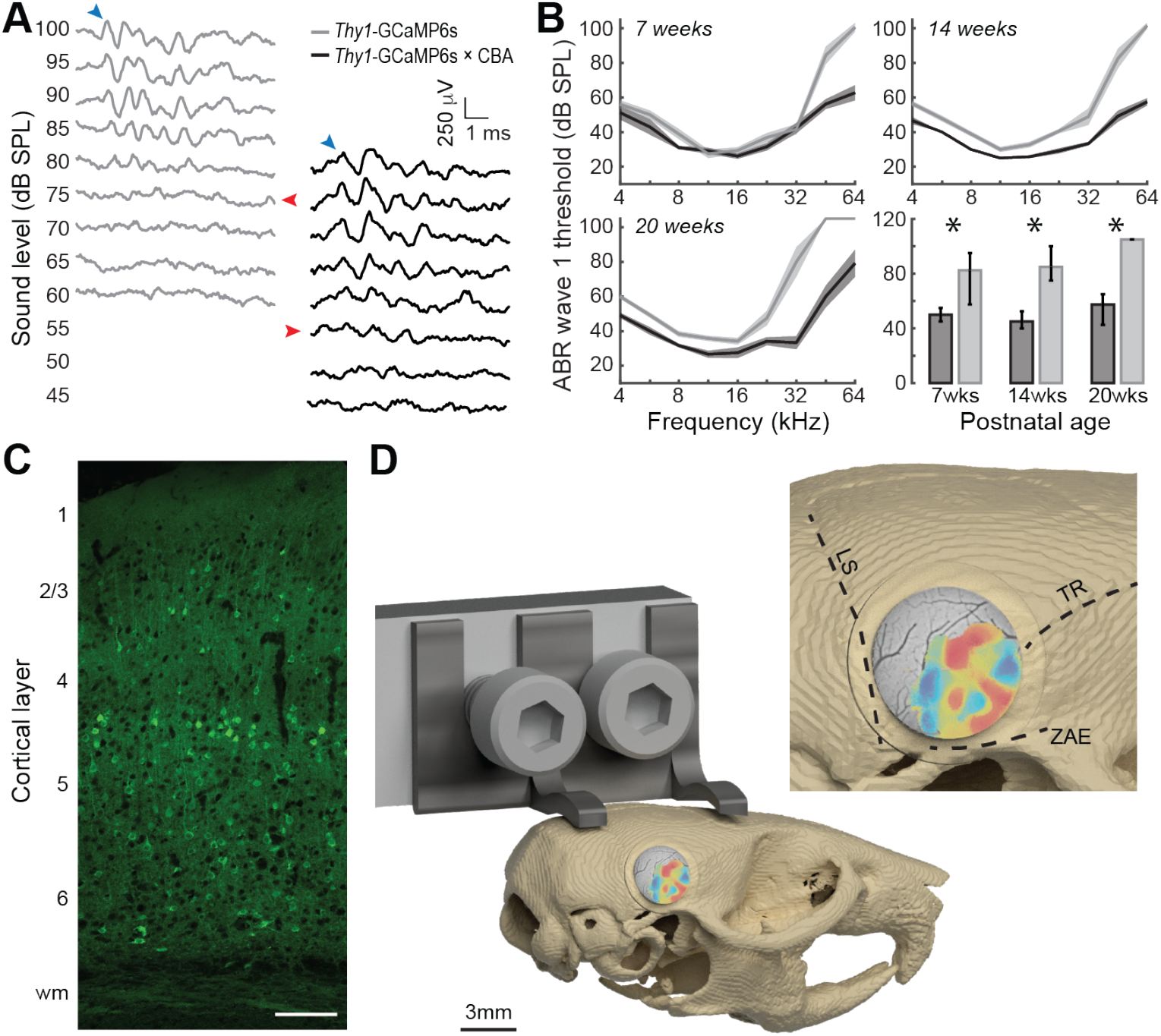
A transgenic GCaMP6s reporter mouse that retains good hearing into adulthood. **(A)** Auditory brainstem response waveforms elicited with 45 kHz tone bursts at various sound levels in a representative 7-week old *Thy1*-GCaMP6s mouse (gray) or the F1 offspring of *Thy1*-GCaMP6s crossed with CBA (black). Blue arrows denote wave 1. Red arrows point towards wave 1 threshold sound level. **(B)** Mean ± SEM ABR thresholds measured at various postnatal ages from *Thy1*-GCaMP6s (n = 27 ears from 14 mice) and *Thy1*-GCaMP6s × CBA mice (n = 23 ears from 12 mice). Bottom right sub-panel presents median and 90% confidence intervals for high-frequency thresholds (32-64 kHz). Asterisks denote p < 0.05 with Wilcoxon Rank-Sum test. **(C)** Confocal image of ACtx GCaMP6s labeling across cortical layers in the *Thy1*-GCaMP6s × CBA mouse. **(D)** Rendering of custom head fixation hardware and placement of the chronic cranial window on mouse skull. *Inset:* Cranial landmarks used to position the window atop the ACtx include the lambdoid suture (LS), temporal ridge (TR) and zygomatic arch extension (ZAE).

We performed widefield imaging in awake, head-restrained *Thy1*-GCaMP6s × CBA mice using a tandem lens epifluorescence microscope that provided a large (5.6mm × 5.6mm) field of view (**Fig. 2A**). We focused the microscope 0.2mm below the pial surface to de-emphasize contributions from surface blood vessels. However, widefield calcium signals collected with epifluorescence microscopes are an amalgam of many cortical layers, not just layer 2/3. Sound-evoked Ca^2+^ signals were temporally de-trended (**Fig. 2B**), filtered (**Fig. 2C**) and pure tone response thresholds were independently calculated for each pixel (**Fig. 2D**). Frequency response areas (FRA) were calculated for each pixel and a best frequency (BF) was computed from the weighted sum of responses at levels ranging from threshold to 20 dB above threshold (**Fig 2E**). The ACtx was defined from the contiguous region of tone-responsive pixels, revealing a well-defined pattern of tonotopically organized gradients of BF that closely resembled prior microelectrode mapping and widefield calcium imaging datasets (Guo et al. 2012; Issa et al. 2014; Joachimsthaler et al. 2014; Liu et al. 2019).

**Figure 2.**
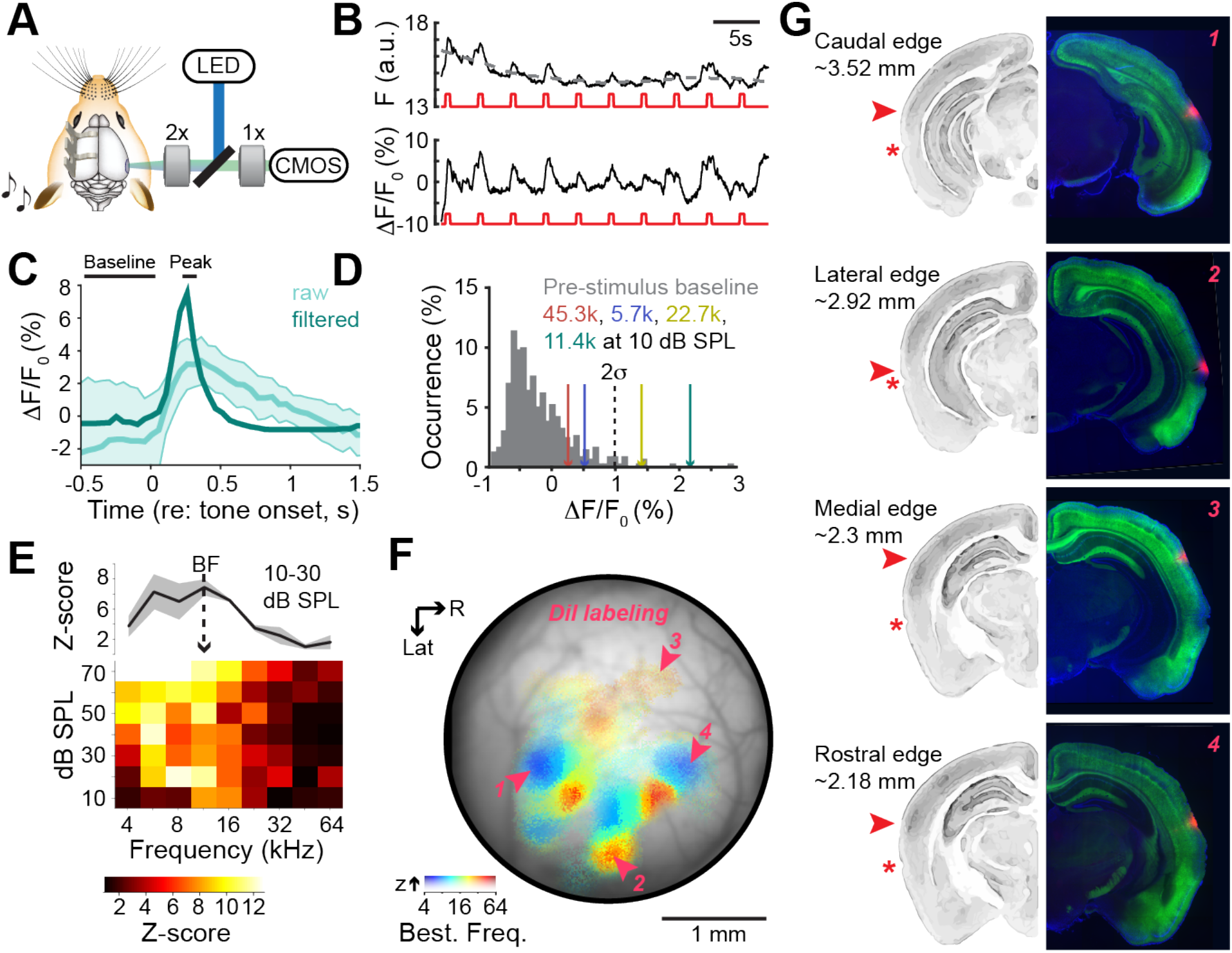
Anatomical landmarks for tonotopically organized fields in the mouse auditory cortex. **(A)** Tandem lens macroscope for widefield GCaMP6s imaging. **(B)** *Top:* Raw calcium signals from a typical pixel are de-trended with a 10s moving average (gray dashed line). *Bottom:* Fractional change in fluorescence is computed relative to the moving average (F_0_). Red lines denote timing of individual tone bursts. **(C)** Mean ± SEM fractional change in fluorescence from 20 repetitions of a 60 dB SPL tone before and after spatial and temporal filtering (light and dark green, respectively). **(D)** Histogram of fractional change values from all temporal baseline periods (gray) and from the peak amplitude of the tone-evoked responses at four frequencies. Dashed black line denotes 2 SD above the mean of the baseline values used to define tone threshold for each pixel. **(E)** Frequency response area of peak responses expressed as z-score from the baseline distribution. Best frequency (BF) is computed from the frequency response function derived from sound levels at threshold to threshold + 20 dB. **(F)** Each pixel is assigned an opacity and hue to denote the response amplitude and tone frequency corresponding to the BF, respectively. In this example case, the medial, lateral, caudal and rostral edges of the tonotopically organized areas were marked with Di-I after imaging. **(G)** Coronal sections of the four Di-I placements from shown in *F* (right) and a grayscale image from a generic mouse brain database to show approximate anatomical landmarks (left). Values express approximate distance from Bregma. Asterisk denotes rhinal sulcus. Red arrows denote center of Di-I expression. Data from panels B-E and panels F-G come from two different mice.

In a subset of mice, the cranial window was removed at the conclusion of imaging and an electrode coated in Di-I was inserted into the most caudal, dorsal, rostral and lateral edge of the tonotopically organized map (**Fig. 2F**). Di-I labeling in ACtx areas with strong tonotopic organization covered a larger area than might be expected from widely used mouse brain atlases. For instance, the lateral edge of the tonotopic zone lies just above the rhinal fissure in a region labeled as Ectorhinal cortex or temporal association cortex (TeA) in the Paxinos and Allen Brain atlases (**Fig. 2G**, second row). Further, the most medial edge corresponds to an area labeled secondary auditory cortex, dorsal area (AuD; **Fig. 2G**, third row).

### Parceling the fields of mouse auditory cortex

Our widefield imaging data confirm the known arrangement of low-frequency hubs at the caudal and rostral edges of the ACtx with tonotopic gradients that branch out and collide with one another to form boundaries between individual fields (**Fig. 3A**). We did not observe a tone-insensitive region at the border of A1 and AAF (Issa et al. 2014, 2016; Liu et al. 2019). Although individual pixel thresholds and response amplitudes can be weaker in this area, we observed a clear low-high-low BF gradient across A1 and AAF, as would be expected from the mirror reversal between core fields documented in over 20 species (Kaas 2011). The tonotopic gradient of A1 was more akin to a fan radiating from its caudal low-frequency hub, than a single linear tonotopic gradient. In the rat, the ventral limb of this gradient is called the ventral auditory field, based on careful characterization of specialized selectivity for low-intensity sounds and separate anatomical inputs from the medial tonotopic limb (Wu et al. 2006; Polley et al. 2007; Storace et al. 2010, 2011). A few new features were apparent that had gone unnoticed in prior microelectrode mapping studies, presumably on account of their very lateral position (**Fig. 3B**). We noted a second low-frequency area at the caudal-lateral extreme of the ACtx map that appeared homologous to an area described in rat imaging experiments as the ventral posterior auditory field (VPAF) (Kalatsky et al. 2005). In addition, we noted a high-frequency area at the lateral extreme of A2 that was more extensive than noted in previous publications.

**Figure 3.**
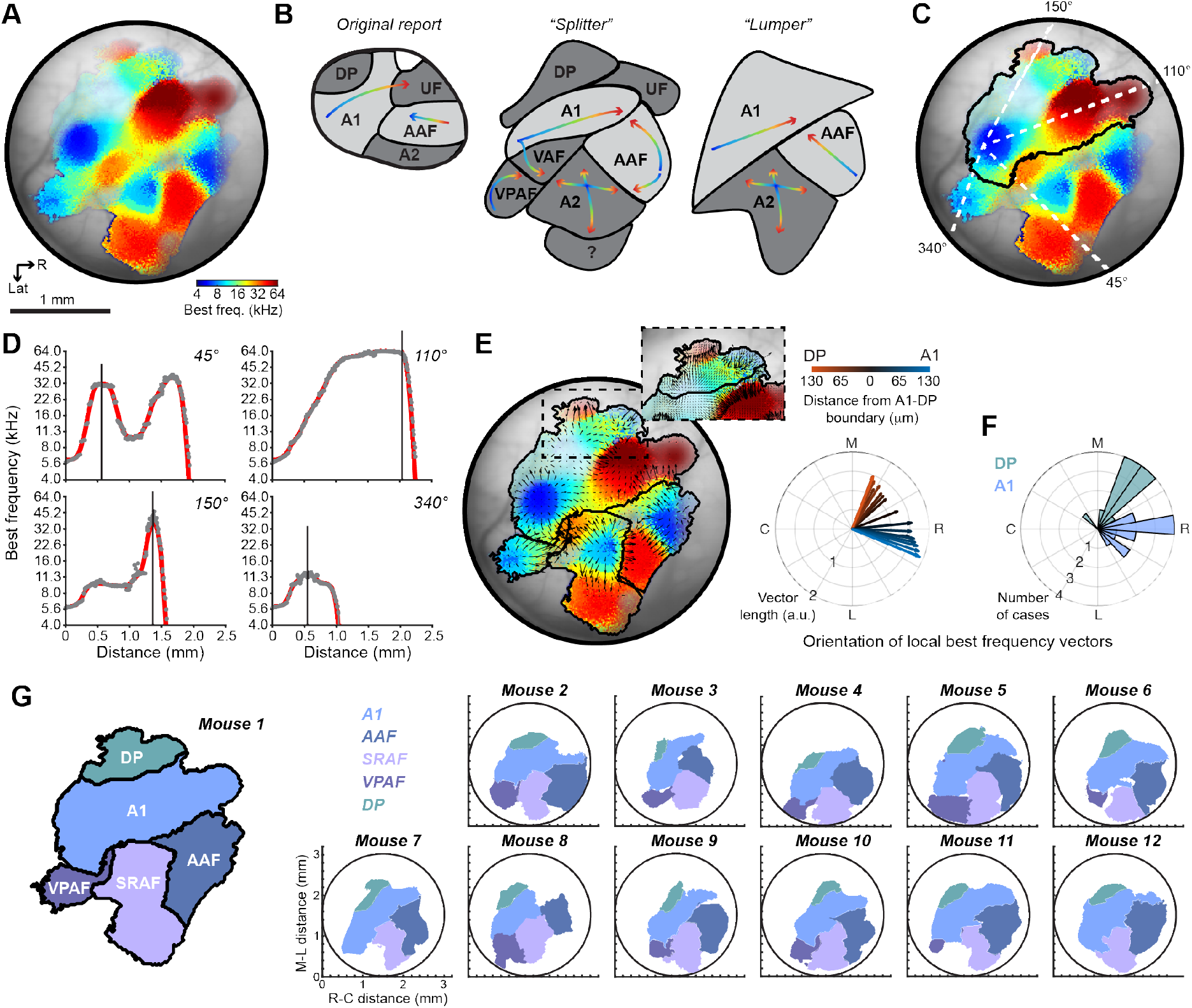
Data-driven parcellation of auditory cortical fields. **(A)** BF map from an example mouse. **(B)** Field boundaries can be established qualitatively, based on reversals or shifts in BF. Field outlines differ from the original characterization of mouse ACtx fields due to the imaging of more lateral brain areas, but boundary positions vary widely depending on “lumper” vs “splitter” biases. **(C)** BF changes along four vectors stretching out from the low-frequency hub in A1. **(D)** Boundaries (vertical lines) are placed at reversals or steep drops in signal amplitude. **(E)** Tonotopic vector map of BFs with boundaries established around each low frequency hub. *Inset:* An abrupt shift in the phase of local BF vectors is used to demarcate the boundary of the dorsal-posterior field. Divergent tonotopy at the boundary of A1 and DP can be appreciated from the progressive shift in BF vector angles with distance from the DP-A1 border. **(F)** Histogram of mean A1 and DP vector angles within 0.13mm of the boundary for each individual case. **(G)** Resultant parcellation of five fields: dorsal posterior (DP), primary auditory cortex (A1), anterior auditory field (AAF), ventral posterior auditory field (VPAF) and the suprarhinal auditory field (SRAF). Maps of the five auditory fields are shown for the same example mouse (left) and the remaining eleven mice used in this study.

The traditional approach for parceling cortical fields is subjective, where bespoke boundaries are drawn at points of reversals or abrupt shifts in the tonotopic gradient. This process is not straightforward in very small cortices, where “separate” gradients can measure less than 100 microns. In these cases, demarcating one field as separate from another can reflect the psychological disposition of the experimenter as a “lumper” or a “splitter”, as much as reflecting any degree of biological ground truth (**Fig. 3B**). These observations motivated us to develop an objective approach for parceling the mouse cortex into separate fields. We projected 360 radial vectors emanating from the center of each of four low-frequency hubs located in A1, AAF, VPAF and A2. Looking at four example vectors from the A1 low-frequency zone, we plotted the BF for each pixel along the vector projection and noted points where the BF reversed or the signal dropped below threshold as boundary points (**Fig. 3C-D**). We then computed a vector map onto each of the four fields, where the length and orientation of the arrows reflect the magnitude and direction of the local BF gradients (**Fig. 3E**). The clear bifurcation of the BF gradient phase along the medial edge of A1 was used to demarcate the boundary of the dorsal posterior field (DP; **Fig. 3E, inset**), where local BF vectors became increasingly divergent with increasing distance from the A1-DP boundary (**Fig. 3E, polar plot**). Non-overlapping distributions of mean vector angle calculated near the A1-DP boundary (± 0.13mm of the boundary) in each individual mouse confirmed the tonotopic separation of A1 and DP (**Fig. 3F**).

This approach settles on a parcellation scheme intermediate to boundaries drawn with a lumping or splitting bias. These data support the position and orientation of A1, AAF and DP, as described in the original mouse electrode mapping study (Stiebler et al. 1997) and confirmed by many subsequent studies (Linden et al. 2003; Hackett et al. 2011; Guo et al. 2012; Issa et al. 2014; Joachimsthaler et al. 2014; Shepard et al. 2015). As we have argued previously, the ultrasonic field (UF) is a misnomer as there is no reason to suggest this area is anything apart from the continuous high frequency gradient of A1, as confirmed from the continuum of BF changes from the central nucleus of the mouse inferior colliculus (Garcia-Lazaro et al. 2015). We further argue that A2 was incorrectly identified in the seminal mouse ACtx mapping study as a homologue to the secondary auditory field found in cats (Reale and Imig 1980; Schreiner and Cynader 1984). Physiologically guided iontophoretic injections of retrograde tracers identify MGBv as the predominant source of thalamic input to A2, not higher-order thalamic sub-divisions ((Ohga et al. 2018), but see (Ji et al. 2016)). Further, A2 units show vigorous responses to pure tone bursts with receptive fields organized into a coarse tonotopic gradient (Guo et al. 2012; Issa et al. 2014). Absent compelling evidence that A2 receives its predominant source of input from the higher-order thalamus or exhibits any functional feature consistent with a higher-order cortical area, it seemed most prudent to name it based on its anatomical position. In rats, the most closely-related evolutionary model system to mice, the auditory field located just medial to the rhinal sulcus is widely referred to as the suprarhinal auditory field (SRAF) (Polley et al. 2007). We now refer to this field as SRAF in mice as well. Although the size and position of each auditory field varies from mouse to mouse, the overall gestalt is preserved in all cases studies here (**Fig. 3G**).

### Analysis of maps and modules in the mouse auditory cortex

BF is the only response feature that is mapped along the extent of each field as a smoothly varying gradient (**Fig. 4A**). We quantified the strength of tonotopy by plotting the local BF phase vectors within each field from a single mouse (**Fig. 4B**, thin gray lines) and calculating the vector sum (**Fig. 4B**, thick black line). Vector strength is derived from the length of the vector sum and reflects the consistency and strength of the local BF gradients. We computed the tonotopic vector strength from a single imaging session in each mouse (N=12) and compared differences across the five fields. To estimate the tonotopic strength that would occur by chance, the BF assignment for each pixel within a field was randomized before calculating the vector strength. This process was repeated 10,000 times and the results were averaged. We observed that the tonotopically organized vector strength from the actual maps was significantly greater than the shuffled maps for all five cortical fields (Wilcoxon signed-rank tests, p < 0.005 for all fields; **Fig. 4C**). Although all fields of the mouse cortex were tonotopically organized, the strength varied between fields (listed in order of strongest to weakest: A1, AAF, SRAF, VPAF and DP; ANOVA, F = 10.49, p < 0.000001). We found that the strength of tonotopic organization in A1 was significantly greater than SRAF, VPAF and DP, but was not significantly different than AAF (post-hoc pairwise comparisons, p < 0.005 after Holm-Bonferroni correction for multiple comparison; A1 vs AAF, p = 0.33), in agreement with our prior electrophysiological mapping study (Guo et al. 2012). We confirmed that the strength of tonotopy in A1 is significantly reduced when BF is calculated at a single, suprathreshold level (70 dB SPL) rather than near threshold (Fig. 4C, post-hoc pairwise comparisons, p = 0.04 after Holm-Bonferroni correction for multiple comparison). Differences between near-threshold and suprathreshold tonotopy trended in the same direction for other fields but were not statistically significant, probably on account of being underpowered for multiple comparisons (post-hoc pairwise comparisons, p > 0.09; Fig. 4C solid versus open bars).

**Figure 4.**
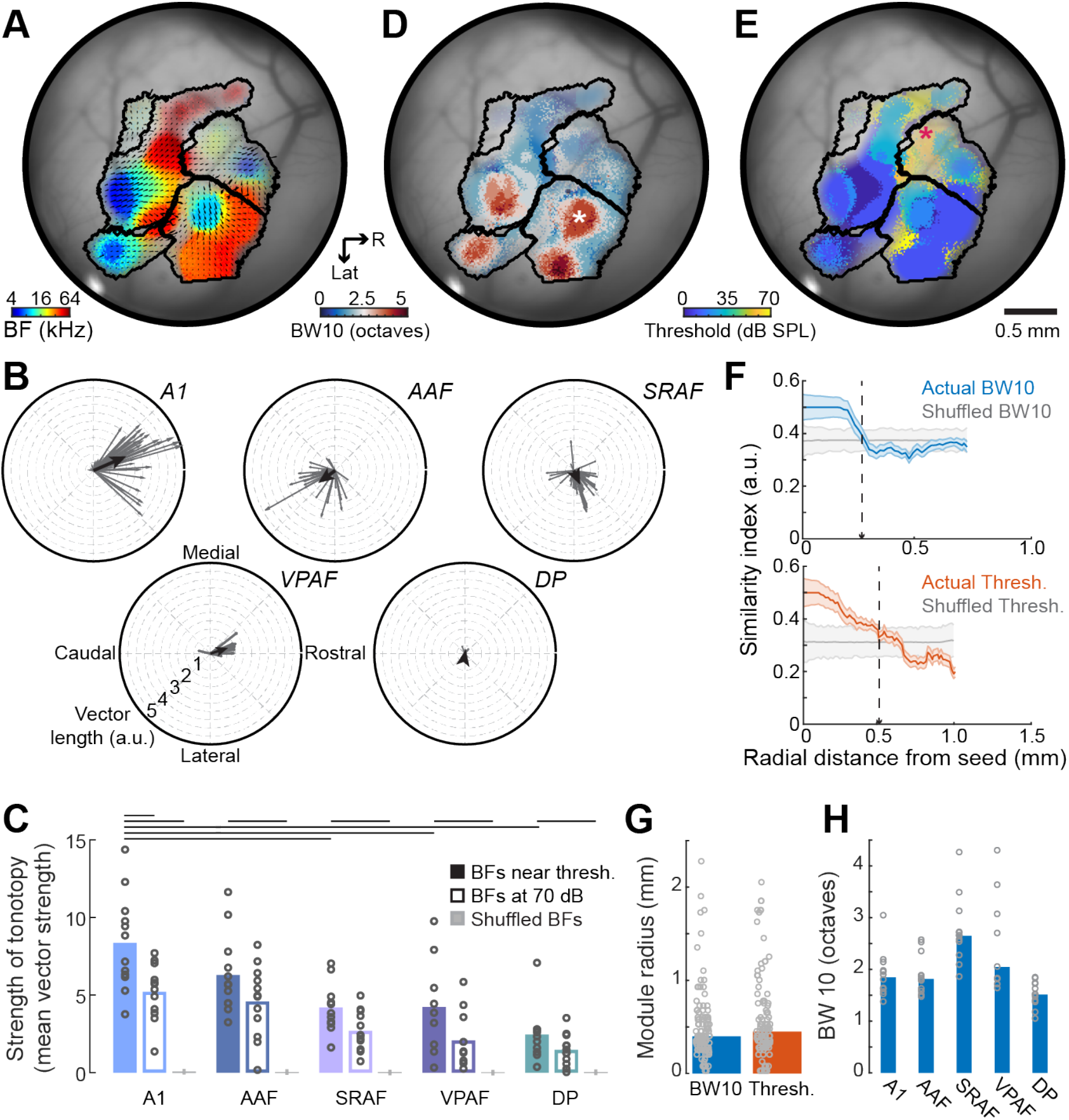
Maps and modules in the auditory cortex. **(A)** BF map from another example mouse where superimposed arrows denote the direction and strength of local BF vectors for each field. **(B)** Distribution of individual BF vectors shown in *A* grouped according to field. Black lines indicate the average vectors. **(C)** Mean (black bars) and individual mouse (gray circles) tonotopic vector strength from the actual BF maps derived from sound levels near threshold (solid bars), at 70 dB SPL (open bars) and shuffled maps (gray bars). Lines indicate statistically significant differences with Wilcoxon Sign Rank tests after correcting for multiple comparisons. Each point in the BF map can alternatively be color coded for frequency tuning bandwidth **(D)** or minimum response threshold **(E). (F)** Mean ± SEM BW10 (top) and threshold (bottom) module size from regions identified with asterisks in *D* and *E* are determined by computing similarity for all pixels relative to center of identified region in the actual and spatially shuffled (gray) maps. Module boundary is marked where actual similarity first overlaps with the similarity that would occur by chance (vertical dashed lines). **(G)** Radial distance for BW10 and threshold modules averaged across all cases (bars) or shown for each individual module (gray circles). **(H)** BW10 for each field averaged across all mice (blue bar) or shown for all individual mice (gray circles).

Other than BF, each of the pixels in the mouse ACtx can also be assigned a value based on tone-evoked response features such as tuning bandwidth or response threshold. As described in cats and rats, the ACtx features circumscribed modules with broad pure tone selectivity (**Fig. 4D**) or high response thresholds (**Fig. 4E**) (Schreiner and Mendelson 1990; Recanzone et al. 1999; Cheung et al. 2001; Read et al. 2001; Polley et al. 2007). A seed was positioned at the center of each individual bandwidth or threshold module and calculated the similarity of the corresponding response feature for all pixels along radial vectors fanning out in 360° from the seed. The mean similarity index across all vectors was then plotted as a function of radial distance and compared to the similarity that would occur by chance in maps where the pixels have been spatially scrambled (**Fig. 4F**, colored versus gray lines). The radial length of individual tuning modules was approximately 0.4mm for both BW10 and threshold (unpaired t-test, t = −1.08, p = 0.83; **Fig. 4G**), indicating that a few separate modules containing regions with homogenously narrow or broad frequency tuning could fall within the boundaries of a single cortical field. In terms of overall differences between fields, we confirmed prior reports that frequency tuning bandwidth was greater overall in SRAF than in A1, AAF and DP, but not VPAF (Wilcoxon rank-sum test, p < 0.05 for each contrast after correction for multiple comparisons, p = 0.5 for SRAF vs VPAF; **Fig. 4H**).

### Long-term stability of tonotopic maps

The chronic cranial window preparation provided long-term optical access to the ACtx, allowing us to measure the stability of frequency tuning over time. Imaging data from an example mouse conveys that macroscopic features are fairly stable over a 37-day period, while the BF of individual pixels suggests some modest session-to-session variability (**Fig. 5A**). We formalized this by registering the widefield images collected from any individual mouse and then computing the absolute value of the BF difference for all pixels that maintained frequency tuning for any given pair of imaging sessions. On average, we found that the BF for any given pixel varied by approximately 0.4 octaves between imaging sessions, which could reflect the true variability in underlying frequency tuning as well as measurement error due to image registration, threshold estimation, internal state variation, photobleaching, window clarity and myriad other experimental factors (**Fig. 5B, top**). To calculate what the BF difference would be by chance, we shuffled the pixels within each field and repeated the measurement. The average BF difference in the shuffled control decreased for smaller fields with reduced ranges of BFs, but was significantly greater than the actual BF difference in all fields (paired t-tests, t > −5.0, p < 0.000005 for each comparison). BF variability did not differ between fields (ANOVA, F = 0.37, p = 0.83) and did not systematically change as a function of interval between imaging session for any field (linear relationship between session and BF difference, p > 0.14 for all fields; **Fig. 5B, bottom**). These findings suggest that the frequency selectivity within a local patch of ACtx (1 pixel equals ~13 μm) varies by less than a half octave over time, confirming prior reports of relative stability of adult cortical sensory maps in the absence of perturbations of sensory experience or afferent activity levels (Jenkins et al. 1990; Masino and Frostig 1996; Polley et al. 2004).

**Figure 5.**
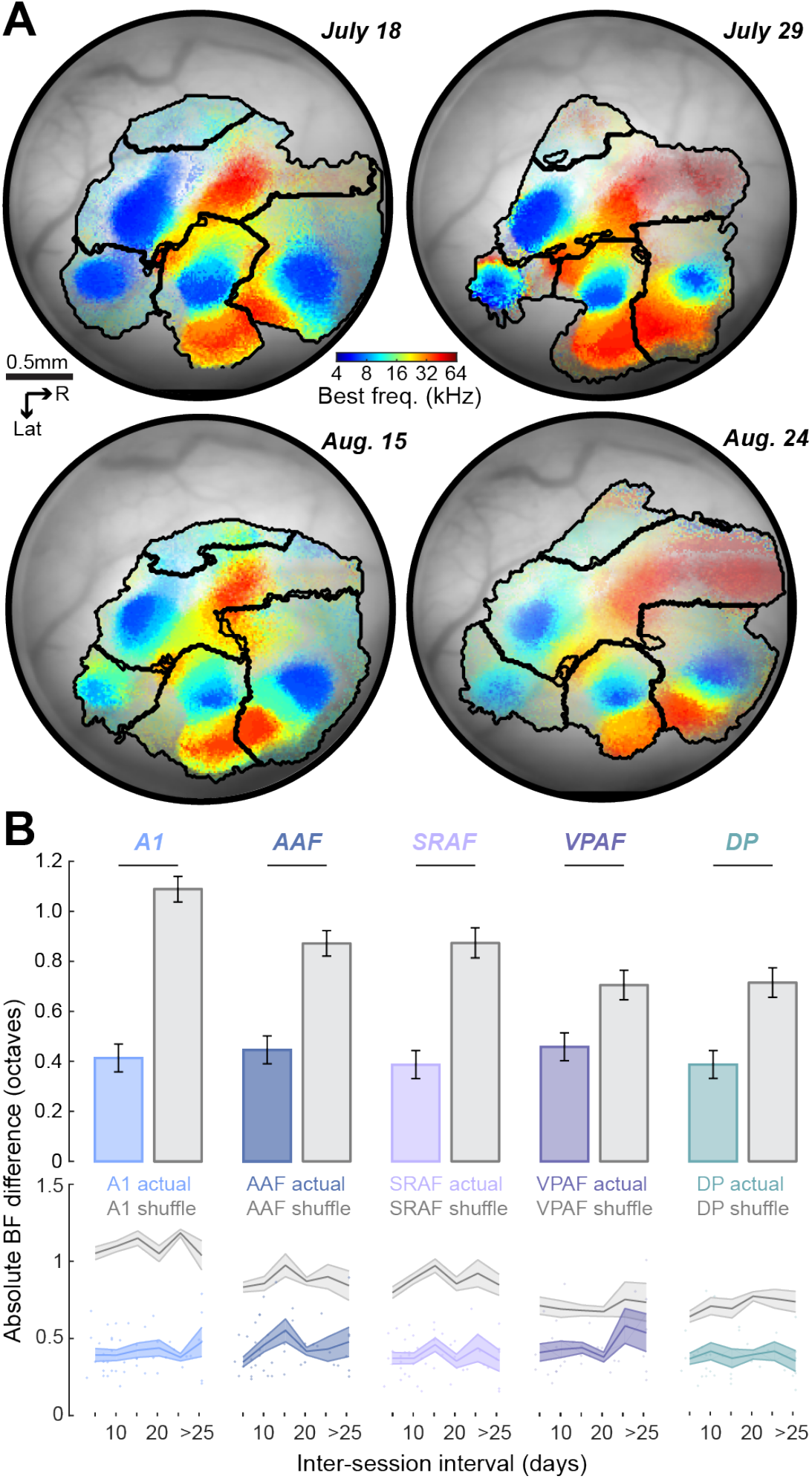
Tonotopic map stability over time. **(A)** Registered BF maps with field boundaries from the same mouse imaged four times over a 37d period. **(B)** Absolute value of the BF difference computed for the same tone-responsive pixels in any pair of images. *Top*: Mean ± SEM of the BF difference from the actual maps (colored) or shuffled maps (gray). Horizontal lines indicate statistically significant differences with a Wilcoxon Sign Rank test. *Bottom*: Mean ± SEM of BF difference as a function of the inter-imaging interval. Individual data points are represented as circles.

### Multiscale imaging of auditory cortex

Whereas the mesoscale tonotopic organization of core fields in mouse ACtx has been confirmed time and again with microelectrode mapping, intrinsic signal imaging or widefield calcium imaging, the underlying organization at a cellular scale remains a point of dispute. The seminal ACtx 2-photon calcium imaging studies relied on bulk-loaded calcium dyes in anesthetized mice and reported that BFs between neighboring L2/3 cells often varied by an octave or more and only weakly conformed to a global tonotopic gradient (Bandyopadhyay et al. 2010; Rothschild et al. 2010). A subsequent study using genetically encoded calcium sensors in awake mice reported a striking precision of local frequency tuning, where the BFs of individual neurons were virtually perfectly aligned to the global tonotopic gradient (Issa et al. 2014). Recent GCaMP6 imaging studies in the ACtx of awake mice suggest that the BFs of neighboring neurons are homogeneous than what was reported in the seminal studies, although some local scatter can be qualitatively appreciated from their example images (Kato et al. 2016; Kuchibhotla et al. 2017).

We reasoned that two factors could affect the correspondence between cellular and mesoscale measurements of ACtx frequency tuning: First, we noted that the study reporting homogeneous local BF tuning did not analytically remove the influence of neuropil from the fluorescence signals measured around individual L2/3 soma (Issa et al. 2014). We expected that fluorescence arising from the surrounding axons and dendrites would reflect the aggregate frequency tuning of the local cellular neighborhood, would more closely match the bulk widefield Ca2+ signal, and should produce more homogenous local BFs. Second, unit recordings and imaging studies have observed that some L2/3 neurons are driven by tones but have poorly defined, irregular FRAs that cannot be accurately described with a singular BF. Our prior study applied the d-prime statistic (d’) to neural FRAs and concluded that the tonotopic organization in L2/3 was substantially degraded when neurons with low d’ were included (Guo et al. 2012).

We expected that including neuropil fluorescence and selecting cells with well-defined FRAs would favor homogenous local BFs that closely matched global tonotopic BF gradients. Conversely, removing neuropil contamination and computing a BF for all tone-responsive neurons independent of tuning quality would produce more heterogeneous local BFs with a coarse global tonotopic organization. To test these predictions, we performed 2-photon imaging of L2/3 pyramidal neurons from a cohort of mice that had undergone widefield imaging 3-14 days prior (N=4, **Fig. 6A**). We spatially registered the fields of view from the tandem lens widefield microscope to the 2-photon microscope so that the tuning of individual neurons could be directly matched to the surrounding mesoscale tonotopic gradient (**Fig. 6B**). With the imaging fields aligned, we quantified frequency tuning for individual neurons before and after calcium signals from the surrounding neuropil were analytically removed from individual neural somata. Somatic FRAs with and without neuropil correction could then be compared directly to signals derived from the corresponding set of pixels from the widefield map. While some L2/3 cells showed robust frequency tuning (**Fig. 6C**), many other cells showed patchy, discontinuous frequency response areas (**Fig. 6D**). The strength of frequency tuning could be quantified by the d-prime statistic (d’), which was used to compute the statistical separability between responses near the BF versus frequency-intensity combinations far away from the BF. Clearly defined frequency tuning (d’ > 1) was only observed in approximately half of L2/3 cells that had significant tone-evoked responses and this fraction was significantly lower in VPAF (n = 587 neurons) compared to A1 (n = 1,482), AAF (n = 2,163) and SRAF (n = 1416; Wilcoxon rank sum tests, corrected with Holm-Bonferroni, p < 1 × 10^−6^ for each comparison; **Fig. 6E**).

**Figure 6.**
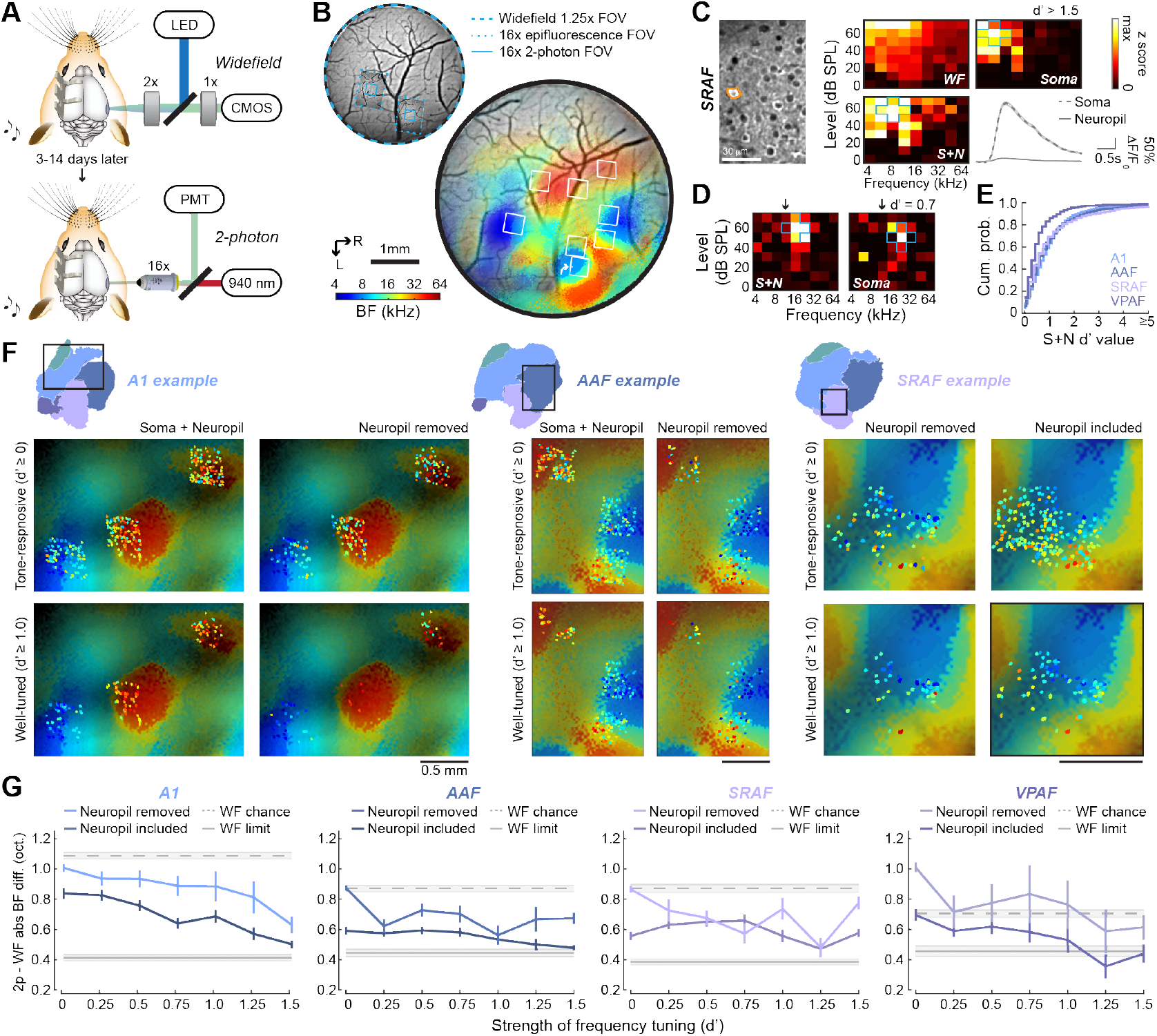
Spatial alignment of mesoscale and cellular frequency tuning. **(A)** Schematic of widefield and 2-photon imaging systems. **(B)** *Top:* Field of view (FOV) registration between the widefield and 2-photon imaging systems. *Bottom:* BF widefield map from an example mouse with 2-photon imaging FOVs superimposed. **(C)** A cell (orange outline) identified with an arrow in *B* is shown with 2-photon excitation. The frequency response area are derived from the z-scores of 2-p GCaMP6s signals after neuropil correction (soma) before neuropil correction (S+N) and from the matching pixels in the widefield system (WF). Mean ± SEM of the fractional change in fluorescence identified by the analysis software arising from the soma or from neuropil. **(D)** An additional cell recorded in with weaker, patchy tuning for frequency and lower d’ values. Blue outlines in *6D-E* identify the five reference frequency-level points used for the calculation of d’. Downward arrows indicate the BF of the corresponding pixels from the widefield image. **(E)** Cumulative density histograms of d’ values in each cortical field. **(F)** Widefield BF maps are extracted from rectangular regions of interest identified in three different mice. Widefield tonotopy is presented in the background with superimposed 2-photon BFs from individual cells. Cellular BF is shown before (left) versus after (right) neuropil correction and for all sound-responsive neurons (top) versus only neurons with FRA d’ values greater than 1.0 (bottom). **(G)** Absolute value of the BF difference between individual neurons and the underlying widefield map are shown for four cortical areas as a function of frequency tuning strength. Dashed gray line represents the mean ± SEM of the BF difference that would occur by chance. Solid gray line represents the mean ± SEM of the smallest possible BF difference, defined from the difference between widefield vs widefield imaging sessions shown in Figure 5.

As predicted, cellular BFs in A1, AAF and SRAF more closely match the global tonotopic maps when comparisons are limited to cells with clearly defined frequency tuning (d’ > 1) and when the local neuropil contribution is not removed (**Fig. 6F**). To formalize these observations, we compared the absolute difference in BF between individual neurons to the corresponding pixels of the widefield BF map. We first defined the lower and upper bounds of the BF difference range from the chronic widefield imaging dataset reported above (Fig. 5). We reasoned that the difference between an individual neuron and the widefield map recorded in two different imaging sessions would not likely be smaller than an individual widefield pixel compared to itself across two imaging sessions (WF limit, solid gray line **Fig. 6G**). Conversely, the BF difference occurring by chance can be defined by the widefield BF difference between a given reference pixel and a randomly selected comparison pixel from a second imaging session (WF chance, dashed gray line **Fig. 6G**).

We analyzed frequency tuning from tone-responsive L2/3 neurons in four mice using the same criteria and analysis methods as the widefield imaging data. We confirmed that BF tuning from individual neurons was a significantly better match to the widefield map before the neuropil contribution is removed (Kruskal-Wallis for A1, AAF and VPAF, p < 0.01; SRAF, p = 0.054). After neuropil correction, the BF difference between individual neurons and the widefield signal was significantly reduced for neurons with stronger overall frequency selectivity (Kruskal-Wallis, p < 0.000001 for all fields; Fig 6F). For neurons with poor frequency selectivity, the alignment to the widefield BF gradient is close to chance, consistent with an underlying heterogenous salt and pepper organization (Bandyopadhyay et al. 2010; Rothschild et al. 2010). However, for neurons with well-defined pure tone receptive fields, the alignment to the widefield map is significantly stronger and approaches the measurement limit (approximately 0.4 octaves), in agreement with 2-photon imaging of genetically encoded calcium indicators in awake mice (Issa et al. 2014, 2016). By accounting for the influence of neuropil and frequency tuning strength, our data suggest that the two ostensibly contradictory descriptions of mouse ACtx organization – locally heterogenous or locally ordered – might, to a degree, both be correct.

### Local and global organization of tonotopy at a cellular scale

As a final step, we explicitly analyzed local BF heterogeneity from our 2-photon imaging data that had been corrected for neuropil contamination and focused on the observation that the distribution of local BFs could be substantially greater when all tone-responsive neurons are included (**Fig. 7A, top**) than when only well-tuned neurons are considered (**Fig. 7A, bottom**). To quantify local BF scatter, we applied a d’ threshold to each field of view to include all tone-responsive neurons (d’ ≥ 0) or to be restricted to neurons with increasingly high d’ values. For each d’ threshold, we treated each cell as a reference and identified all other cells within a 50 μm radial distance. We computed the median BF across all cells within a given local neighborhood and determined the BF difference for each cell relative to the median BF value (Tischbirek et al. 2019). We iterated this process for each neuron in the field of view and compiled a histogram of BF differences, using the interquartile range of the BF difference distribution as an index of local BF heterogeneity (**Fig. 7C**). Local BF scatter was significantly dependent on the strength of frequency tuning in A1, AAF and SRAF, but not in VPAF (permutation tests, p < 0.005 for A1, AAF and SRAF, p = 0.64 for VPAF, respectively; **Fig. 7D**). This analysis confirms that in the core fields, estimates of local BF scatter could vary by a factor of two based solely on the inclusion criteria for selecting candidate neurons for analysis. In VPAF, by contrast, the degree of scatter is independent of frequency tuning quality and appears inherently more variable than other cortical fields.

**Figure 7.**
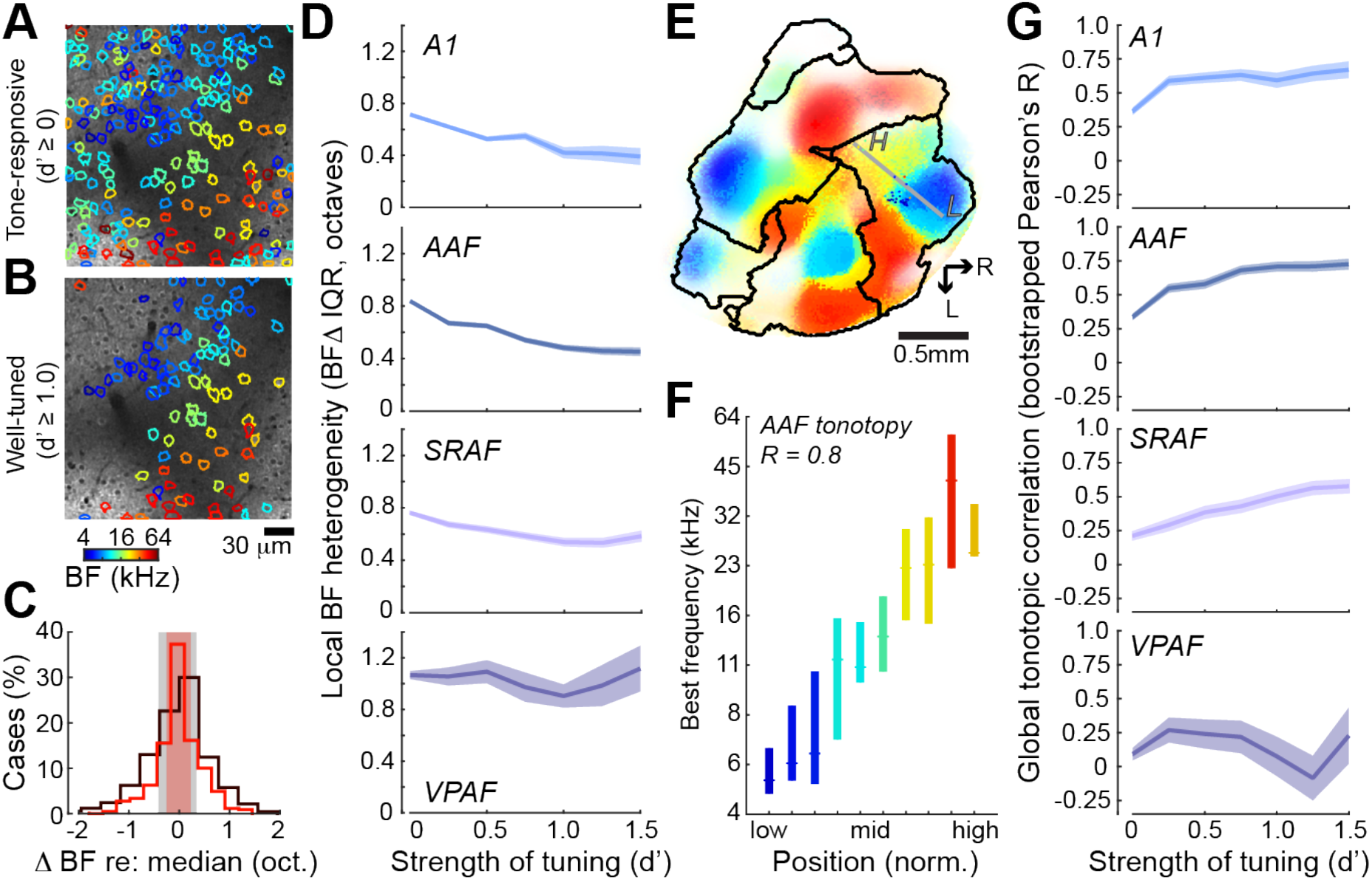
Local and global organization of tonotopy. **(A-B)** 2-photon FOV showing cellular BFs from all sound-responsive neurons versus neurons with strong frequency tuning (d’ > 1, *B*). For a given reference cell, all neighboring cells within a 50 μm radius are identified. The median BF for all cells within this local neighborhood is computed. **(C)** The difference in BF for each cell versus the neighborhood median is shown for all sound-responsive neurons (black) or for neurons that are strongly tuned to frequency (d’ > 1, red). The interquartile range of the BF difference histogram (shaded regions) provides an estimate of local BF heterogeneity. **(D)** Mean ± SEM BF heterogeneity as a function of frequency tuning strength for four cortical areas. **(E)** Widefield BF map from an example mouse with superimposed 2-photon cellular tuning from two FOVs in AAF. Gray line indicated global tonotopic vector. L = low. H = high. **(F)** Individual cells across AAF FOVs in all mice are assigned to a position in the global tonotopic vector. Median (horizontal line) and interquartile range of BFs within each AAF position bin. The linear relationship between the mesoscale tonotopic position and cellular BF is indexed by the Pearson’s R value. **(G)** Mean ± SEM of the bootstrapped Pearson’s R as a function of frequency tuning strength for four cortical areas.

As a corollary to local BF heterogeneity, we quantified the linear relationship between the BF of individual cells and their position along the low-to-high frequency extent of the corresponding field. Reports emphasizing local BF heterogeneity have described a very coarse correlation to global tonotopic position (Pearson R ~ 0.2), while reports of homogeneous local BFs describe a precise linear correlation (Pearson R ~ 0.9) (Bandyopadhyay et al. 2010; Rothschild et al. 2010; Issa et al. 2014). To address these observations in our data, we projected the position of each individual cell onto a vector that connected the low and high end of the global tonotopic gradient for the corresponding field (**Fig. 7E-F**). We then computed the Pearson correlation coefficient between BF and tonotopic position. As suggested from our prior analyses, the global correlation strength, operationally defined here as the bootstrapped Pearson correlation coefficient, could be accurately described as coarse or precise, depending on the inclusion criteria for allowing tone-responsive neurons into the analysis. Global correlation strength was significantly greater for neurons with higher d’ values in A1, AAF and SRAF, though again not in VPAF, where cellular tuning did not conform to an overall tonotopic scheme (Permutation test, p < 0.001 and p = 0.77, respectively; **Fig. 7G**).

## Discussion

In this study, we performed multiscale imaging from a transgenic mouse that expressed a genetically encoded calcium indicator in excitatory neurons throughout the cerebral cortex but retained excellent hearing into adulthood (**Fig. 1**). We described a procedure to pre-process the widefield epifluorescence signals and identify an auditory response threshold and BF for each pixel individually (**Fig. 2**). We marked the caudal, rostral, medial and lateral extremes of the ACtx with Di-I and noted that the lateral edge of the tonotopically organized map extended farther caudally and laterally than would be expected from the anatomical landmarks identified in widely used mouse brain atlases. We implemented a data-driven parcellation approach that used abrupt drops in signal strength and reversals or phase shifts in BF gradients to identify five cortical fields – A1, AAF, SRAF, VPAF and DP (**Fig. 3**). We observed statistically significant mesoscale tonotopic organization in all fields, where modules of similar frequency tuning bandwidths and response thresholds were superimposed on BF gradients (**Fig. 4**). Frequency tuning remained relatively stable over approximately one month of repeated imaging from the same mice, where the BF of individual pixels varied by less than 0.5 octaves (**Fig. 5**). We used 2-photon imaging to spatially register the frequency tuning of individual neurons to the widefield maps. We found that correspondence between the cellular and mesoscale tonotopic maps ranged from slightly better than chance to nearly equal to the measurement limit, depending on the strength of cellular frequency tuning and whether neuropil contributions were factored out or included (**Fig. 6**). With the neuropil contamination removed, we found that the degree of local BF scatter and orderly progression of local BFs along the tonotopic axis both reflected the strength of frequency tuning (**Fig. 7**).

### Precision of frequency tuning in the mouse auditory cortex

Tonotopy is among the most rudimentary aspects of ACtx organization and is clearly predictive of frequency guided auditory behaviors (Jenkins and Merzenich 1984; Znamenskiy and Zador 2013). Yet, the ostensibly straightforward question of whether the ACtx is tonotopically organized has remained a point of contention for over fifty years (for review see (Kanold et al. 2014)). Opposing papers have debated whether the tonotopic gradient in core fields was coarse or precise from the start of the modern neurophysiological era (Evans and Whitfield 1964; Merzenich and Brugge 1973) and continuing right up to the current era (Bandyopadhyay et al. 2010; Rothschild et al. 2010; Issa et al. 2014). A consensus view is emerging in the rodent somatosensory and visual cortex for a globally ordered topographic mapping of the receptor epithelium comprised of locally heterogeneous tuning at a cellular scale (Ohki et al. 2005; Sato et al. 2007; Bonin et al. 2011; Clancy et al. 2015). In ACtx, there is clear support for a robust tonotopic organization in the core auditory fields when maps are made at mesoscale resolution, where individual points of measurement reflect a pooling of local activity. Whether studied with microelectrode multiunit recordings from the middle layers of anesthetized animals (Hackett et al. 2011), widefield imaging of intrinsic signals (Moczulska et al. 2013; Kato et al. 2016; Tischbirek et al. 2019) or genetically encoded calcium sensors from the upper layers of awake animals (here and in (Issa et al. 2014; Liu et al. 2019)), a highly ordered systematic progression of preferred frequency is readily apparent in the core fields of the mouse ACtx, in keeping with descriptions in over 20 other mammalian species (Kaas 2011).

Zooming into any local cellular neighborhood within the tonotopic map reveals a considerable heterogeneity of preferred frequency between neighboring single neurons, as revealed by single unit spiking of neighboring units (South and Weinberger 1995) or suprathreshold calcium events recorded with 2-photon imaging (Bandyopadhyay et al. 2010; Rothschild et al. 2010; Winkowski and Kanold 2013; Liu et al. 2019). The balance of globally ordered, smoothly changing gradients built from heterogeneous, inherently ‘noisy’ local processing is consistent with reports of globally ordered topographic mapping of MGB thalamocortical projections (Hackett et al. 2011), yet heterogeneous frequency tuning within individual MGB axon terminals (Vasquez-Lopez et al. 2017) or between individual spines of L2/3 pyramidal neurons (Chen, Leischner, et al. 2012).

The only remaining point of debate is centered on the degree of local heterogeneity in preferred frequency tuning. On one end of the spectrum, 2-photon imaging from layer 2/3 neurons in awake mice that express GCaMP3 suggested extremely low variance in preferred frequency between local neurons in A1 (standard deviation of 0.4 octaves for a 230×230 μm field of view) and a tight correspondence between local preferred frequency and the global tonotopic gradient (correlation coefficient > 0.85) (Issa et al. 2014). However, this report did not analytically correct for neuropil contribution. Even in visually isolated soma, calcium signals recorded with 2-photon microscopes average fluorescence across tens of microns in the z-axis, where much of this signal will reflect a contribution from out-of-focus axons and dendrites from other neurons. As we show here, including the neuropil substantially reduces local BF heterogeneity and improves the alignment to the global tonotopic map (Fig. 6). Therefore, we would argue that this study may have over-estimated the degree of local BF precision by including a more global neuropil signal (Lee et al. 2017). On the other end of the spectrum, the original pioneering 2-photon imaging studies of mouse ACtx described substantial heterogeneity from neuropil-corrected L2/3 cells in anesthetized animals. Increased heterogeneity is unlikely to arise solely from the anesthetized state, as systematic differences in BF tuning are not observed in cells recorded in both anesthetized and awake conditions (Guo et al. 2012; Tischbirek et al. 2019). Some of the heterogeneity reported in these studies could reflect the use of bulk-loaded dyes, which would increase the contribution of fluorescence signals from non-neuronal cell types and certain upper layer GABAergic interneurons that may have broader frequency selectivity than pyramidal neurons (Li et al. 2014).

Another possibility is that these seminal studies either used tone bursts with very short inter-trial intervals (0.4s, (Rothschild et al. 2010)) or longer (1s duration) amplitude-modulated tones (Bandyopadhyay et al. 2010). Recording from neurons in an adapted state or from neurons responding during sustained periods of stimulus-related activity could increase the complexity of auditory tuning and increase the heterogeneity between neighboring neurons (South and Weinberger 1995; Wang et al. 2005; Kato et al. 2017) or may conflate the independent frequency tuning for sound onset versus offset (Liu et al. 2019).

The degree of local heterogeneity reported here is somewhere in between these two extremes and is qualitatively consistent with our prior descriptions from electrophysiological recordings of single units (Guo et al. 2012) and a recent demonstration of cellular tuning across all layers of the A1 column (Tischbirek et al. 2019). The most important point from the data presented here was that correspondence to the widefield map (Fig. 6G), the degree of local BF heterogeneity (Fig. 7D) and the global correlation (Fig. 7G) could differ by a factor of two, depending on the inclusion of neurons that were activated by tones, but were poorly selective for a single, narrow range of frequencies. This is completely self-evident and corresponds exactly to the same observation we made previously in electrophysiological recordings from L2/3 units (Guo et al. 2012). Essentially, if a neuron had an irregular, broad or multi-peaked receptive field, its frequency preference cannot be as accurately reduced to a single number. Regardless, even by limiting our sample of neurons to those with reasonably strong frequency selectivity, it was clear that the BFs of neighboring neurons *i)* do not reflect a salt and pepper organization, but rather are strongly predicted by their position within the overlying mesoscale map, but *ii)* vary on the order of approximately half an octave in any local neighborhood (~50 μm XY radial distance). This degree of local heterogeneity could be an unavoidable consequence in small brains with substantial divergence of thalamocortical and intracortical connectivity (Hackett et al. 2011). Reduced heterogeneity would be expected along a radial column than along the tangential plane, or in species with larger auditory cortices or more precise anatomical connectivity (Atencio and Schreiner 2010, 2013; Guo et al. 2012; See et al. 2018; Tischbirek et al. 2019), but we expect to see that globally systematic yet locally heterogenous selectivity would be an organizing feature of sensory cortex organization (Kanold et al. 2014).

### Organization and naming schemes for multiple fields of the mouse auditory cortex

Whereas there is general agreement about the balance between global order and local diversity in A1 and AAF, there is no consensus on what fields of the mouse ACtx are “higher-order”, where they are located or even what they should be called. Here, we implanted a cranial window to cover the full extent of the mouse ACtx, affording us optical access to lateral areas of the cortex that are difficult to record from with acutely inserted microelectrodes (**Fig. 8A**). Looking across individual mice (**Fig. 8B**), we consistently observe four low-frequency hubs at the edges of the ACtx that fan out and collide with one another to form the boundaries between fields (**Fig. 8C, left**). In keeping with the seminal mapping study as well as the nomenclature adopted in other species, the objective parcellation approach used here is consistent with having A1, AAF and DP labeled as separate fields (**Fig. 8C, right**). The frequency gradients identified by our mesoscale GCaMP6 imaging are a close fit to that described by earlier mapping of GCaMP3 signals with one exception: They argue that there is a tone-insensitive region at the border of A1 and AAF (**Fig. 8D**, (Issa et al. 2014, 2016)). We confirmed that pixels in this area have higher response thresholds, but with the individual pixel thresholding procedure used here it is evident that even though tone-driven responses are weaker in this region, they have BFs that are consistent with the overall frequency gradients linking A1 and AAF, in keeping with over 20 other mammalian species and with prior microelectrode mapping of the A1-AAF junction in the mouse (Linden et al. 2003; Hackett et al. 2011; Kaas 2011; Guo et al. 2012; Shepard et al. 2015). Prior calcium imaging (**Fig. 8D**) and microelectrode mapping studies also identified an area with low-frequency BFs lateral to A1 and AAF (**Fig. 8E** and **8F**, respectively (Stiebler et al. 1997; Guo et al. 2012; Issa et al. 2014; Joachimsthaler et al. 2014; Ohga et al. 2018)). By developing a surgical approach to position the cranial window more laterally and caudally than previous studies, we identified a fourth low-frequency hub that we named VPAF, in keeping with a description of a similar ventral-posterior field identified with widefield imaging of intrinsic signals in the rat ACtx (Kalatsky et al. 2005).

**Figure 8.**
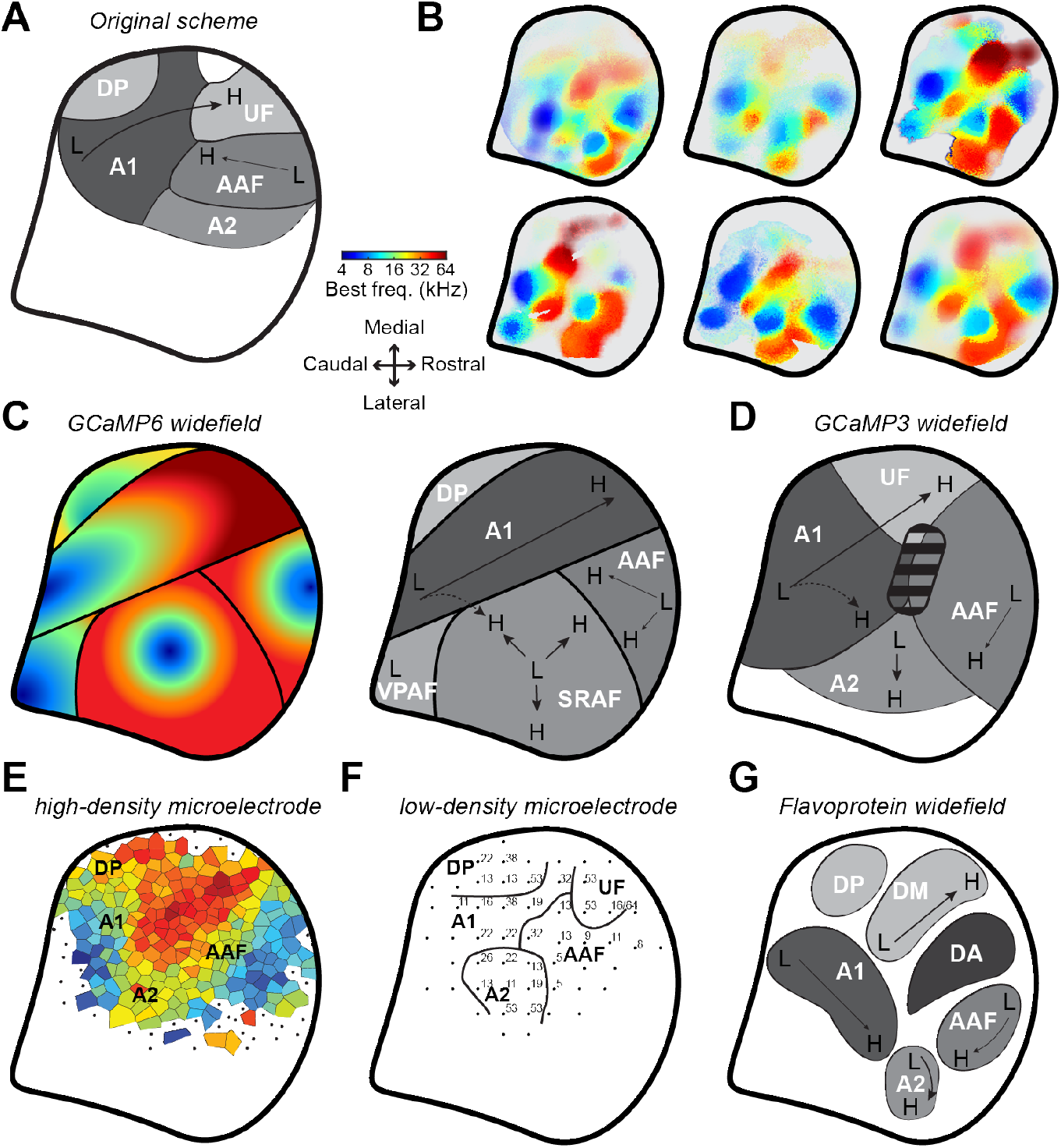
Mapping the core and higher-order fields of the mouse auditory cortex. **(A)** Schematic of the original proposal for the layout of the mouse ACtx fields (Stiebler et al. 1997). Thick black line represents the areal outline of the complete ACtx studied here. **(B)** BF maps from six individual mice. **(C)** *Left:* Cartoon of the general BF gradients and field boundaries suggested from the GCaMP6 imaging data reported here. *Right:* low-to-high frequency gradients (L and H, respectively) and field naming designations proposed here. **(D)** Low-to-high frequency gradients and field naming designations proposed from earlier GCaMP3 widefield imaging studies (Issa et al. 2014, 2016). Striped region denotes a tone-insensitive region identified by these studies. **(E)** Voronoi tessellation of preferred frequency and field naming designations from a high-density multiunit microelectrode mapping study (Guo et al. 2012). **(F)** Preferred frequency and field naming designation identified from a lower density microelectrode mapping study (Joachimsthaler et al. 2014). **(G)** Cortical parcellation scheme suggested from flavoprotein widefield imaging studies (Tsukano et al. 2015, 2016; Tsukano, Horie, Takahashi, et al. 2017). Note that all schematics have been adapted, re-annotated and resized from their original form to fit the right hemisphere.

Although naming conventions vary, the overall tonotopic gestalt is consistent across these widefield calcium imaging and microelectrode mapping studies. The only exception comes from a series of reports using widefield imaging of endogenous flavoprotein signals that describe a low-frequency area interposed between A1 and AAF and mis-identify the upper limb of A1 as belonging to a separate field referred to as DM (Tsukano et al. 2015, 2016; Tsukano, Horie, Ohga, et al. 2017; Tsukano, Horie, Takahashi, et al. 2017) (**Fig. 8G**). Flavoprotein fluorescence signals are an order of magnitude slower and weaker than genetically encoded calcium sensors. Possibly on account of the need for longer trial durations, these studies generally test a more limited set of tone frequencies at a single sound level, as compared to the 72 frequency/level combinations used here. As tonotopy is substantially degraded when derived from tones presented at a single suprathreshold sound level (Fig. 4C and (Guo et al. 2012), the organizational scheme suggested from these studies should be interpreted cautiously.

Mouse ACtx researchers have generally adopted the original naming scheme proposed by the seminal low-density microelectrode mapping studies of the mouse ACtx (Fig. 8A and 8F). We propose that some aspects of the original scheme are misleading and should be changed in favor of a naming system that is more consistent with auditory cortical fields in other mammals. The designation of an ultrasonic field (UF) should be abolished on the grounds that *i)* “ultrasonic” is anthropomorphic and refers to any frequency above the limit of human hearing (approximately 20 kHz) and could therefore refer to approximately half of the mouse hearing range and not just the representation of frequencies above 45 kHz; *ii)* there doesn’t appear to be anything discrete or discontinuous about the cortical representation of frequencies above 45 kHz; in most cases it simply appears as the high frequency elaboration of the A1 and AAF tonotopic gradients, in keeping with the elaboration of high-frequency BFs described in the tonotopic representations of the mouse central nucleus of the inferior colliculus (Garcia-Lazaro et al. 2015).

Further, we argue that A2 is a misnomer for the field described here as the suprarhinal auditory field. There was never a particularly compelling reason to label this area as A2 to begin with. The seminal study relied on low-density microelectrode mapping to note the frequency tuning bandwidths were wider and the tonotopic organization less clear than in the core fields (Stiebler et al. 1997), both of which have been confirmed here (Fig. 4). The designation of a secondary auditory field is only widely used in cats but is most comparable to a non-tonotopically organized parabelt areas in non-human primates (Reale and Imig 1980; Schreiner and Cynader 1984). In other rodents, carnivores and nonhuman primates, field naming conventions follow the anatomical position of the field and not its presumed position in a hierarchy of cortical processing. Among the commonly used animal models for ACtx research, laboratory rats (*Rattus*) are the closest evolutionary relative to the laboratory mouse (Mus). Because the suprarhinal auditory field described here shares the same position and tonotopic orientation as the rat, we argue that this field should also be called SRAF (Polley et al. 2007). Perhaps more importantly, higher-order auditory fields – by definition – receive their predominant thalamic input from higher-order thalamic subdivisions (e.g., MGBd), not primary thalamic subdivisions (MGBv) (Rose and Woolsey 1949a, 1949b; Andersen et al. 1980; Winer et al. 2005). Dual neuranatomical tracer injections into A1 and SRAF in the mouse revealed that both fields receive inputs from separate zones of the MGBv, with hardly any input from MGBd (Ohga et al. 2018). Although a second study that did not reconstruct the full rostral-caudal extent of the MGB and did not use physiological guidance for their tracer injections came to the opposite conclusion (Ji et al. 2016), at a minimum the source of thalamic input to these areas is uncertain and deserves further study with physiologically guided tracer injections before any attribution of a primary or secondary level of processing can be made.

### Where – if anywhere – are the higher order fields of mouse auditory cortex?

Unlike the visual cortex, where hierarchies for stimulus processing abound, differences in the nature and form of auditory stimulus processing between fields of the ACtx are more a difference of degree than a difference of kind. Three notable exceptions have been identified. First, strictly non-primary areas of the ACtx have been identified in the human brain, where lateral regions show specialized responses for music and speech that are not observed in the primary areas (Leaver and Rauschecker 2010; Norman-Haignere et al. 2015; Overath et al. 2015; Kell et al. 2018; Norman-Haignere and McDermott 2018). Second, higher-order areas have been identified in the ferret ACtx that selectively encode sounds according to their behavioral meaning and not their acoustic features (Atiani et al. 2014; Elgueda et al. 2019). Third, a sub-type of neuron with broad spike waveforms have been identified in a higher-order field of songbird ACtx that supports the *de novo* emergence of sparse, contrast-invariant representations of conspecific vocalizations (Schneider and Woolley 2013; Kozlov and Gentner 2016; Ono et al. 2016).

Here, we used pure tone bursts in passively listening mice to delineate the boundaries of cortical fields without revealing much about any underlying specializations. Mesoscale tonotopy was strong in A1, AAF and SRAF (Fig. 4C), where underlying neurons that were well-tuned to sound frequency showed comparably homogeneous BFs (Fig. 7D) and adherence to global tonotopic vectors (Fig. 7G). VPAF and DP exhibited less organized, incomplete representations of the cochlear frequency map. Cellular imaging in VPAF revealed a highly disorganized salt and pepper organization of BFs that was largely insensitive to frequency tuning strength.

Ultimately, differences in sound frequency organization can suggest candidates for higher-order fields, but cannot provide definitive evidence for where a field sits within a cortical hierarchy or heterarchy. “Higher-order” is an anatomical designation that reflects a preponderance of higher-order thalamic inputs, stronger inputs from brain areas that encode multi-sensory inputs, and stronger connectivity with non-auditory structures such as frontal cortex, amygdala or neuromodulatory centers. Functional markers such as tonotopic precision, task-related modulation and cross-modal sensitivity are products of – rather than determinants of – the complement of afferent inputs that define as region as core or higher-order. In this regard, the poorly selective disorganized frequency representations in VPAF and DP suggest a relatively weak input from the MGBv and stronger input from higher-order brain areas, but this can only be demonstrated with carefully positioned injections of tracers or viral vectors. In mouse visual cortex, researchers have made good headway identifying specialized visual feature processing and state-dependent modulation in fields beyond V1 (Glickfeld et al. 2014; Ramesh et al. 2018; Beltramo and Scanziani 2019). In mouse ACtx, researchers have by and large focused on recording only from A1, typically in passively listening animals. Here, we propose that VPAF and DP may represent good candidates for studies on higher-order anatomical connectivity studies as well as neurophysiological experiments that focus on the extraction of auditory features that guide purposeful behavior.

## Acknowledgements

We thank C. Harvey, M. Andermann, G. Yellen, C. Niell, S. Gandhi, N. Afrashteh, R. Froemke and K. Kuchibhotla for sharing advice on surgical protocols, hardware solutions and data preprocessing suggestions for calcium imaging. We thank M. Pachitariu for making Suite2p publicly available. We thank D. Kim and the Genetically-Encoded Neuronal Indicator and Effector (GENIE) Project at the HHMI’s Janelia Farm Research Campus for making the Thy1-GCaMP6s mouse publicly available. SR and AEH prepared the mice and collected data, with additional support from KKC and JR. SR analyzed data, with support from AEH. DBP and AEH designed the experiments. KEH and RSW developed software control. DBP wrote the manuscript, with feedback from all authors.

